# A striatal plasticity that supports the long-term preservation of motor function in Parkinsonian mice

**DOI:** 10.1101/2022.01.10.475638

**Authors:** Joe C. Brague, Rebecca P. Seal

## Abstract

Motor deficits of Parkinson’s disease (PD) such as rigidity, bradykinesia and akinesia result from a progressive loss of nigrostriatal dopamine neurons. No therapies exist that slow their degeneration and the most effective treatments for the motor symptoms: L-dopa -the precursor to dopamine, and deep brain stimulation can produce dyskinesias and are highly invasive, respectively. Hence, alternative strategies targeted to slow the progression or delay the onset of motor symptoms are still highly sought. Here we report the identification of a long-term striatal plasticity mechanism that delays for several months, the onset of motor deficits in a mouse PD model. Specifically, we show that a one-week transient daily elevation of midbrain dopamine neuron activity during depletion preserves the connectivity of direct but not indirect pathway projection neurons. The findings are consistent with the balance theory of striatal output pathways and suggest a novel approach for treating the motor symptoms of PD.

## INTRODUCTION

The dorsal striatum has a critical role in the execution of voluntary movement. GABAergic spiny projection neurons (SPNs), which comprise >95% of striatal neurons, are the major output neurons. The neurons receive dense innervation from both cortical and thalamic excitatory neurons and serve as the primary postsynaptic target of substantia nigra pars compacta (SNpC) dopamine neurons. SPNs in the dorsal striatum form what are referred to as the direct “go” and indirect “no go” pathways (Albin et al., 1989; Lanciego et al., 2012; McGregor and Nelson, 2019; Villalba and Smith, 2010; 2018). Direct pathway SPNs project to the globus pallidus internal and substantia nigra pars reticulata (SNpR) and express the excitatory dopamine 1 receptor (D1R), while indirect pathway SPNs project to the SNpR by way of the globus pallidus external and subthalamic nucleus and express the inhibitory dopamine 2 receptor (D2R). Dopamine acts on both SPN populations to increase the net output of direct over indirect pathways to facilitate movement (Villalba and Smith, 2010).

In Parkinson’s disease (PD), the progressive loss of SNpC dopamine neurons differentially alters the activity of the striatal out pathways, weakening the direct pathway relative to the indirect pathway, and causing progressively more profound motor deficits that manifest as bradykinesia and akinesia (Smith et al., 2009; Stephens et al., 2005). Changes in the activity of SPNs with the loss of striatal dopamine manifest as altered intrinsic electrophysiological properties as well as dendritic morphology and excitatory connectivity. In terms of the intrinsic excitability, in rodent models of PD, loss of dopamine causes excitability of D1R SPNs to increase and of D2R SPNs to decrease (Surmeier, 2007). In terms of morphological changes, postmortem brains of Parkinson’s disease (PD) patients (Stephens et al., 2005; Zaja-Milatovic, 2005), as well as primate (Smith, 2009; Smith and Villalba, 2008; Villalba and Smith, 2010) and rodent models of PD (Day et al., 2006; Fieblinger et al., 2014; Gagnon et al., 2017; Villalba and Smith, 2018) consistently show a marked loss of mature spines and dendritic arbors on both types of SPNs. Interventions that restore the balance of activity within the basal ganglia network, including at the level of the striatal SPNs, are therefore a major focus for correcting the motor deficits in Parkinson’s disease patients.

We previously reported that mice lacking the vesicular glutamate transporter, VGLUT3, do not develop the deficit in forepaw reaching that normally results from dorsal striatal dopamine depletion (Divito et al., 2015). We also reported that the mice exhibit a transient daily hyperdopaminergia that is due to an increase in the synthesis and release of striatal dopamine, and which also manifests in a concomitant increase in locomotor activity (Divito *et al*., 2015). Since the hyperdopaminergia phenotype only occurs during the active cycle, it alone cannot explain the normal motor behavior observed in the knockout (KO) mice following dopamine depletion, which persists throughout the circadian cycle. To determine the mechanistic basis for the normal motor behavior observed in Parkinsonian VGLUT3 KO mice, we therefore used a combination of chemogenetic, pharmacological, histological, and electrophysiological approaches. Here we show that a transient daily elevation of dopamine during dopamine depletion preserves D1R, but not D2R SPN morphology and connectivity and delays the onset of motor deficits for months.

## RESULTS

In addition to the increased release of striatal dopamine and enhanced locomotor activity observed during the active cycle, we also observed a circadian dependent increase in the density of morphologically immature spines on SPNs in VGLUT3 KO mice (Divito *et al*., 2015). These spines were shown to be electrically immature as well. A link between increased striatal dopamine and the formation of immature spines on SPNs has been reported in the context of repeated cocaine administration (Lee and Dong, 2011; Lee et al., 2013). Cocaine enhances striatal dopamine levels and given daily for 5 days, induces the formation of electrically silent synapses on SPNs (Huang et al., 2009). Interestingly, cessation of cocaine administration causes the spines to mature into functional synapses (Lee and Dong, 2011; Lee *et al*., 2013). Given the parallels between the transient daily increase in striatal dopamine that occurs in the KO mice and repeated cocaine administration, we wondered whether the depletion of dorsal striatal dopamine may similarly increase the density of mature spines on SPNs as is observed with cocaine cessation.

We thus measured the densities of mature (mushroom) and immature (long/thin and stubby) spines of SPNs in wildtype (WT) and KO mice using DiI labeling in acute striatal slices. Given that the direct and indirect SPNs have opposing effects on basal ganglia network activity and motor behavior, we specifically marked the direct pathway SPNs by injecting the retrograde tracer cholera toxin b-subunit (CTB)-Alexa-488 into the SNpR *in vivo*. As expected in KO mice at baseline (not depleted), the density of immature spines was increased relative to WT and only during the active cycle. Here we show that this increase occurs on D1R SPNs (WT (black) vs KO (red) p<0.0001, N=3 mice per group, n=7 dendrites per group) (Figure 1A). As has been reported by other groups, in WT mice, dopamine depletion decreased the densities of both spine types on D1R and D2R SPNs regardless of the time of sacrifice (active: baseline vs depleted p<0.0001, N=3 mice per group, n=7-12 dendrites per group; inactive: baseline vs depleted p<0.05, N=3 mice per group, n=6 dendrites per group) (Figures 1A and 1B). In KO mice, on the other hand, mature spines did not decrease but instead were preserved on D1R SPNs after depletion and interestingly, like the behavior, this was observed across the circadian cycle (active: N=3-4 mice per group, n=9-12 per group and inactive: N=3-4 mice per group, n=6 dendrites per group) (Figure 1A). As in depleted WT mice, long/thin and stubby immature spine densities were decreased on both types of SPNs in KO mice throughout the circadian cycle (Active: baseline vs depleted p<0.0001, inactive: baseline vs depleted p<0.000; N=3 mice per group, n=7-12 dendrites per group) (Figures 1A and 1B). As a control, forepaw reaching behavior was measured one week following depletion and just prior to sacrificing the mice. KO mice (N=14) showed the expected normal bilateral forepaw reaching, while WT mice (N=8) developed the ipsilateral paw dominance motor deficit (active: ipsilateral vs contralateral p<0.0001; inactive: ipsilateral vs contralateral p<0.0001) (Figure S1E). We also show that the degree of dopamine depletion, assessed by measuring tyrosine hydroxylase immunoreactivity in the dorsal striatum was similar between KO (N=12) and WT mice (N=14 mice) (Figure S1F). Thus, we report that immature spine density is specifically increased on D1R SPNs in KO mice relative to WT during the active cycle. And remarkably that following dopamine depletion, mature spine density specifically on D1R SPNs is increased in the KO relative to WT mice. Importantly, like the reaching behavior, this preservation of mature spines on D1R SPNs persists throughout the circadian cycle, suggesting the induction of a long-term plasticity.

**Figure 1.**
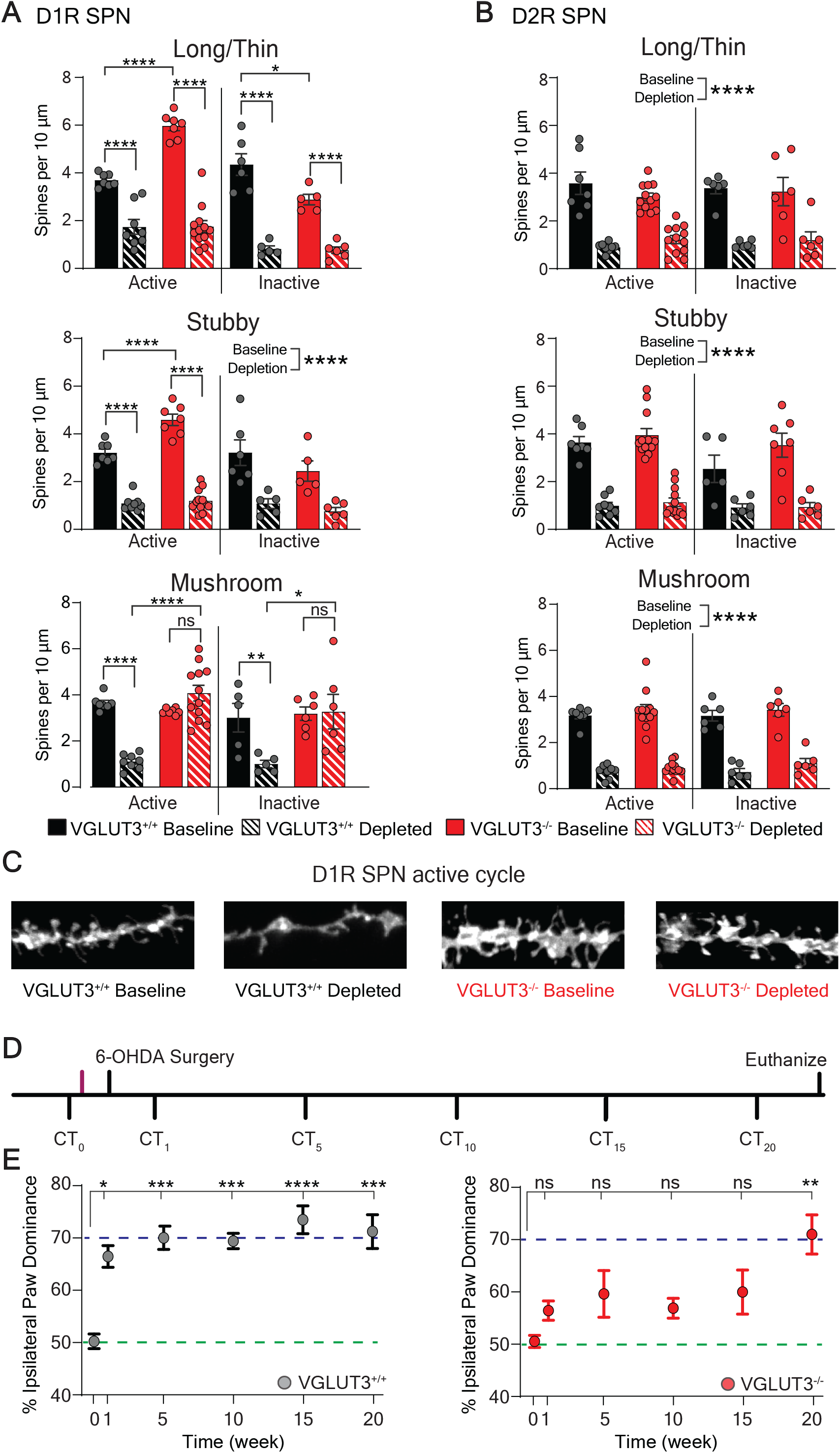
Specificity of spine density phenotypes to D1R SPNs and duration of normal motor behavior in VGLUT3 KO mice. **(A)** At baseline during the active cycle, VGLUT3 KO mice (red solid bar) show enhanced density of immature (long/thin and stubby) spines on D1R SPNs compared to WT controls (black solid bar). After dopamine depletion, mature (mushroom) spine densities are significantly decreased in WT mice (blacked solid vs striped bars) but are preserved in VGLUT3 KO mice (red solid vs striped bars) across the circadian cycle. Densities of all other spine types are decreased significantly after dopamine depletion in both WT and KO mice. Significant interaction by 2-way ANOVA (*genotype* x *condition*) for all spine types and time points (except stubby spines during the inactive cycle; main effect p<0.0001). **(B)** D2R SPN spine densities are similar at baseline in KO and WT mice and all spines are decreased similarly by depletion in both genotypes. Two-way ANOVA (*genotype x condition*), no significant interaction, main effect for condition is reported. **(C)** Representative images of Dil filled D1R SPN dendrites during the active cycle from WT (black) and KO (red) mice. **(D)** Experimental paradigm for measuring duration of preserved motor behavior in KO mice following dopamine depletion. CT = cylinder test. Unilateral dorsal striatal 6-OHDA (6 µg) injection. Magenta line indicates time of spine density measurement at baseline. **(E)** VGLUT3 KO mice (red dots) do not show significant ipsilateral paw dominance until 20 weeks post-depletion. WT mice (gray dots) show ipsilateral dominance at 1 week. Purple and green dotted lines indicate 50% (normal) and 70% (Parkinsonian) ipsilateral paw reaches, respectively. One-way repeated measures ANOVA with Bonferroni post-hoc comparison to baseline *p<0.05, **p<0.01, ***p<0.001, ****p<0.0001, ns = not significant.

Since WT mice clearly develop ipsilateral paw dominance by one week following the injection of 6-OHDA, we chose to measure behavior at this time point in our studies thus far. However, given that the KO shows a behavioral plasticity following dopamine depletion, we continued to measure paw reaching weekly in the KO until they developed the ipsilateral paw dominance. Remarkably, a consistently significant motor deficit was not observed until ∼20 weeks post-depletion (Figures 1D and 1E and Figures S1G and S1H).

### Recapitulation of VGLUT3 KO hyperdopaminergic phenotypes with DREADDs

The preservation of mature spine density selectively on D1R SPNs in VGLUT3 KO mice following dopamine depletion agrees with the hypothesis that we put forth as a parallel to what is observed with cocaine administration and cessation. However, the relationship between our findings and the hyperdopaminergia phenotype is only correlative. Indeed, the plasticity instead could be due to other factors that result from a global loss of VGLUT3. We therefore directly tested whether the transient daily increase in dopamine causes a long-term preservation of mature spine density on D1R SPNs and bilateral forepaw reaching behavior by switching to a mouse model that exclusively recapitulates the hyperdopaminergia phenotype of the KO.

In this model, the excitatory designer receptor exclusively activated by designer drugs (eDREADD), hM3Dq, was targeted to midbrain dopamine neurons (in mice WT for VGLUT3), allowing for the controlled transient elevation of striatal dopamine (Armbruster et al., 2007). An adeno-associated virus expressing Cre-dependent hM3Dq was injected bilaterally into the midbrain of mice expressing Cre recombinase under control of dopamine transporter (DAT) regulatory elements (DAT^Cre^ mice) (Figures 2A and 2B). To match the locomotor activity to what we observe in the KO mice, we injected the ligand for hM3Dq, clozapine-n-oxide (CNO) systemically at 0.1 mg/kg, one hour prior to the onset of the active cycle (Figure 2C). Locomotor activity of these mice was compared to two controls: DAT^Cre^ mice that 1) expressed the mCherry reporter in midbrain dopamine neurons and were injected with 0.1 mg/kg CNO (mCherry+CNO, dark blue) or 2) expressed the eDREADD and were injected with saline (eDREADD+Saline, light blue). The eDREADD mice injected with CNO (referred to hereafter as transiently elevated dopamine, TED, mice) (eDREADD+CNO, orange), showed a significant increase in locomotor activity at the peak of the active cycle compared to the controls (*p<0.05, **P<0.01, ***p<0.001, ****p<0.0001; N=7-11 mice per group) (Figure 2C) and that was similar to the increase in locomotor activity observed with the VGLUT3 KO mice (Figure S1B) (Divito *et al*., 2015).

**Figure 2.**
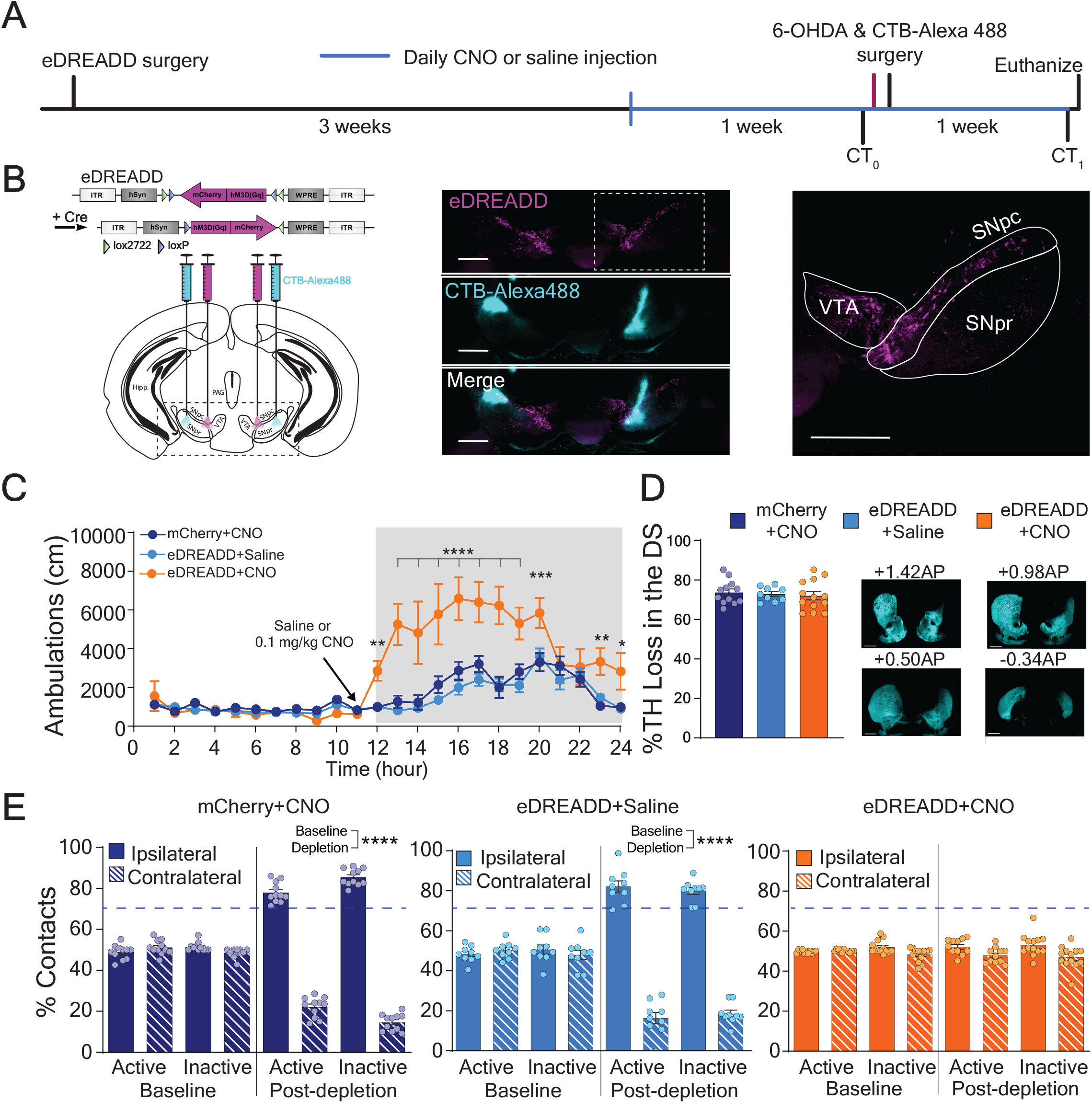
TED mice recapitulate the normal motor behavior observed in VGLUT3 KO mice after unilateral dorsal striatal dopamine depletion. **(A)** Experimental paradigm for transiently elevating dopamine (TED) in the preservation of bilateral paw reaching behavior after dorsal striatal dopamine depletion. CT0= cylinder test at baseline, CT1= cylinder test 1 week after dorsal striatal 6-OHDA injection. Magenta line indicates baseline spine density collection (data shown in Figure 3). **(B)** Left: Illustration of Cre-dependent eDREADD (hM3Dq) viral construct and brain sites for bilateral injection of AAV8 eDREADD and CTB-Alexa488 in DAT^Cre^ mice. Middle: Confocal image eDREADD-mCherry (magenta) in midbrain dopamine neurons and CTB-Alexa488 (cyan) in SNpr to back-label D1R SPNs. Right: Magnified confocal image. VTA-ventral tegmental area; SNpc-substantia nigra pars compacta; SNpr-substantia nigra pars reticulata. Scale bars = 500 µm. **(C)** Locomotor activity is significantly elevated after CNO injection (0.1 mg/kg, i.p.) in eDREADD expressing mice (TED mice, orange line) compared to control mice (dark and light blue lines). Significant interaction by 2-way ANOVA (*genotype* x *condition*). Bonferroni multiple comparison *post-hoc*. **(D)** No difference between TED mice (orange bar) and controls (light and dark blue bars) in percent loss of TH immunoreactivity in the dorsal striatum 1 week after 6-OHDA injection. Kruskal-Wallis test **(E)** TED mice (orange bars) show preserved bilateral paw reaches one week after 6-OHDA injection, while control mice show ipsilateral paw dominance (light and dark bars). No significant interaction by 2-way ANOVA (*paw x condition*) in TED mice. Significant interaction and Bonferroni *post-hoc* analyses for controls. *<0.05, **p<0.01, ***p<0.005, ****p<0.001.

We next tested whether the transient increase in midbrain dopamine neuron activity during the active cycle preserves normal bilateral forepaw reaching in the cylinder one week after unilateral dopamine depletion (Divito *et al*., 2015). Mice were injected with either CNO (0.1 mg/kg, i.p.) or saline one hour prior to the onset of the active cycle for 14 consecutive days while 6-OHDA injected into the dorsal striatum unilaterally on Day 7 (Figures 2A and 2B). Cylinder paw contacts were measured at the peak of the active and inactive cycles one day before and one week after injection of the toxin (Figure 2A). Control mice were tested in parallel. Both lines of control mice showed the expected ipsilateral paw dominance during both active and inactive cycles (mCherry+CNO: ipsilateral vs contralateral, p<0.0001, N=12 mice; eDREADD+Saline: ipsilateral vs contralateral, p<0.0001; N=11 mice) (Figure 2E). In contrast, TED mice showed no paw preference in the cylinder test, which persisted throughout the circadian cycle (N=12-14 mice) (Figure 2E). Importantly, the extent of dopamine depletion in the dorsal striatum was similar between TED mice and controls (Figure 2D). These data provide strong support for the hypothesis that transiently increasing striatal dopamine neuron activity during induction of a Parkinson state produces plasticity supportive of normal bilateral paw reaching behavior.

We next tested whether there were changes in the spine densities of SPNs in TED mice relative to controls at baseline and after dopamine depletion (Figure 3 and Figure S3). At baseline, TED mice showed an increase in the density of long/thin immature spines on D1R SPNs during the active, but not the inactive cycle (Active: eDREADD+Saline vs eDREADD+CNO, p<0.05; N=3-4 mice per group, n=6-9 dendrites per group) (Figure 3A and Figure S3). The D2R SPN spine densities were in contrast similar for all animal groups across the circadian cycle (N=3 mice per group, n=5-9 dendrites per group) (Figure 3B and Figure S3). Following dopamine depletion, mature spine densities on D1R SPNs of control mice but not TED mice were significantly reduced and this was observed across the circadian cycle (active: eDREADD+Saline, baseline vs depleted, p<0.0001; mCherry+CNO, baseline vs depleted p<0.001; N=3-4 mice per group, n=6-7 dendrites per group; eDREADD+CNO, N=3 mice per group, n= 10-11 dendrites per group; inactive: eDREADD+Saline, baseline vs depleted p<0.0001; mCherry+CNO, baseline vs depleted p<0.01; N=3 mice per group, n=5 dendrites per group; eDREADD+CNO, N=3 mice per group, n= 5 dendrites per group) (Figure 3A and Figure S3). The D2R SPN spine densities, in contrast, were decreased significantly across all animal groups throughout the circadian cycle (p<0.0001, N=3 mice per group, n=5-9 dendrites per group) (Figure 3B and Figure S3).

**Figure 3.**
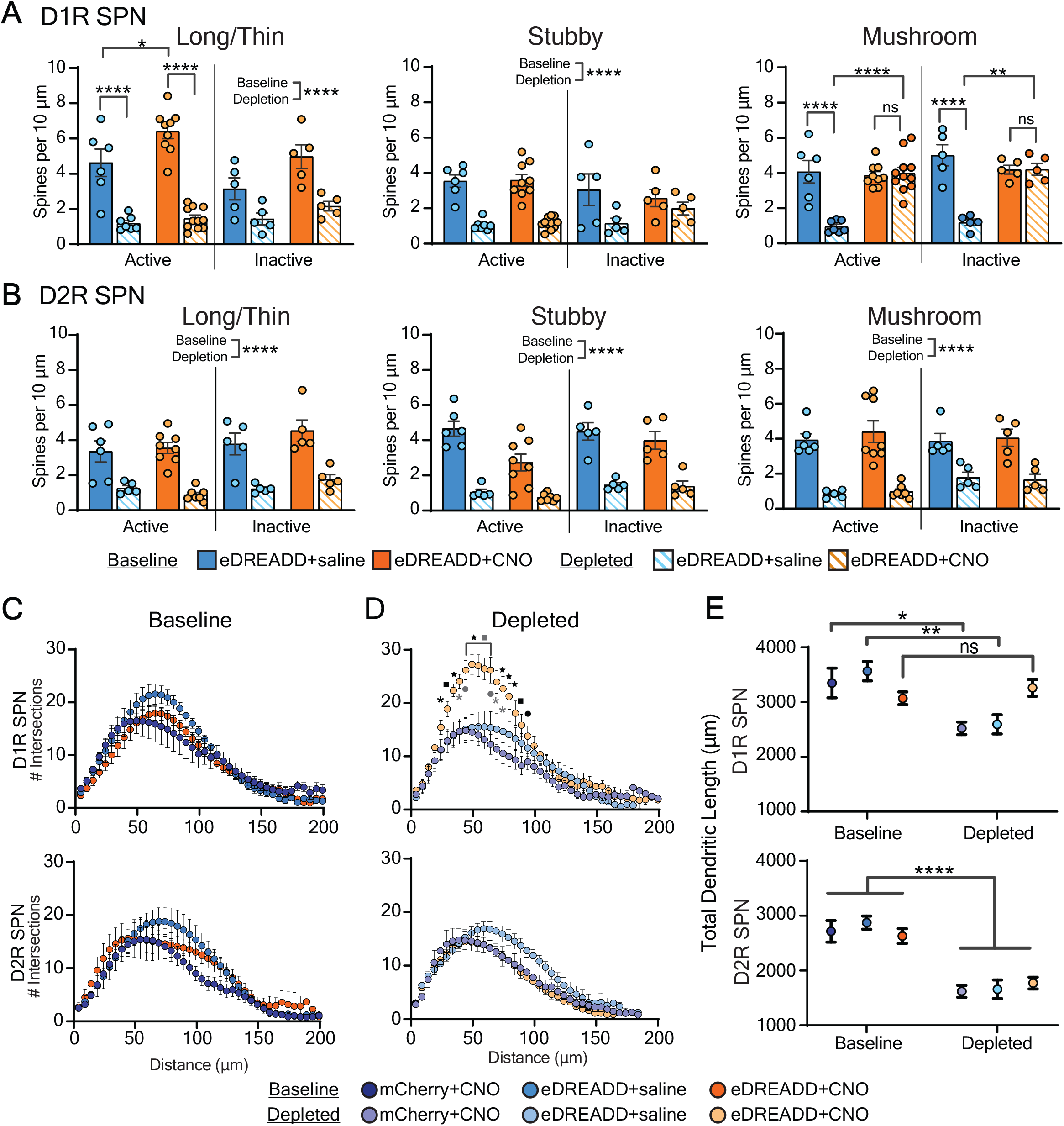
TED mice recapitulate the morphological changes observed in SPNs of VGLUT3 KO mice after dorsal striatal dopamine depletion. **(A)** At baseline, immature (long/thin) D1R SPN spine density in TED mice (orange solid bars) is enhanced compared to control eDREADD+saline (light blue solid bars) only during the active cycle. After depletion, mature (mushroom) spine density is preserved in TED mice (orange solid vs striped bars) but not in control (light blue solid vs striped bars) across the circadian cycle. mCherry+CNO control located in Extended data S3 for illustrative purposes. Significant interaction by 2-way ANOVA (*animal group* x *condition*) mushroom and long/thin spines. Bonferroni post-hoc analyses. No interaction stubby and long/thin spines during inactive cycle. Significant main effect for condition. **(B)** Depletion decreased D2R SPN spine densities to the same extent in TED mice (orange bar) and control (light and dark blue bars). No interaction by 2-way ANOVA (*animal group x condition*). Significant main effect for condition. **(C)** Number of intersections by D1R and D2R SPN dendrites in dorsal striatum is similar across animal groups at baseline (left). No interaction by 2-way ANOVA (*animal group x intersection*). After dopamine depletion, intersections by D1R SPNs in TED mice (light orange dots, right) are significantly enhanced compared to controls (light purple and light blue dots). Significant interaction by 2-way ANOVA (*animal group x intersection*). Bonferroni post-hoc analyses. Black symbols compare Ted mice to mCherry+CNO controls, grey symbols compare Ted mice to eDREADD+Saline controls. *p<0.05; •p<0.01, ▪p<0.001, ★p<0.0001. **(D)** TED mice (dark and light orange dots), but not controls (dark and light purple and blue dots) show preserved total dendritic length of D1R SPNs after dopamine depletion. Significant interaction by 2-way ANOVA (*animal group x condition*). Bonferroni post-hoc analyses. Total dendritic length of D2R SPNs is decreased significantly within all animal groups after depletion. No interaction by 2-way ANOVA (*animal group x condition*). *p<0.05, **p<0.01, ***p<0.005, ****p<0.001, ns = not significant.

Synaptic input is not only a function of mature spine density, but also dendritic arbor length and complexity. We therefore performed Scholl analyses on the SPNs of TED mice and controls (Figures 3C and 3D and Figure S2A). Prior to depletion, the degree of dendritic arbor branching by D1R SPNs of TED mice did not differ significantly from controls (N=3 mice per group, n=5-7 neurons per group) (Figure 3C and Figure S2A). After depletion, however, the extent of branching of D1R SPNs was significantly greater in TED mice than controls (*post-hoc* multiple comparisons: Black symbols compare mCherry+CNO to TED mice, gray symbols compare eDREADD+Saline to TED mice; *p<0.05; •p<0.01, ▪p<0.001, ★p<0.0001; N=3 mice per group, n=5-10 neurons per group) (Figure 3D and Figure S2A). D2R SPN branching did not differ between TED mice and controls before (N=3 mice per group, n=5-7 neurons per group) or after depletion (N=3 mice per group, n=5-10 neurons per group) (Figures 3C and 3D and Figure S2A). In terms of the total dendritic length, the D1R SPNs in control but not TED mice were decreased following depletion (mCherry+CNO, baseline vs depleted p<0.05; eDREADD+Saline, baseline vs depleted p<0.01; N=3-4 mice per group, n=5-11 neurons per group; eDREADD+CNO, N=3 mice per group, n= 9-11 neurons per group) (Figure 3E). The total dendritic lengths of D2R SPNs, in contrast, were significantly decreased after dopamine depletion in all animal groups (p<0.0001; N=3-4 mice per group, n=5-10 neurons per group). Taken together, the findings provide strong causal support for the idea that a transient daily increase in dopamine neuron activity during induction of the Parkinsonian state preserves D1R SPN morphology and forepaw reaching behavior following depletion.

### Intrinsic electrophysiological properties of SPNs after dorsal striatal dopamine depletion

In addition to morphological changes, intrinsic electrophysiological properties of SPNs such as the response to current injection, rheobase, and input resistance also influence excitability and output of SPNs and thus motor dysfunction in models of PD (Fieblinger *et al*., 2014). We therefore characterized the intrinsic properties of SPNs in TED mice and controls, both before and after dopamine depletion. Similar to previous reports (Fieblinger *et al*., 2014), the mean action potential (AP) frequency measured for D1R SPNs was decreased (baseline vs depletion p<0.001, N=5 mice per group, n=5-9 neurons per group), and for D2R SPNs was increased (baseline vs depletion p<0.05, N=5 mice per group, n=5-9 neurons per group) and this was observed for all animal groups (Figure 2B and Figure 2C). Additionally, the resting membrane potential (Vrest) of D2R SPNs was significantly more negative for all groups (p<0.01, N=3 mice per group, n=5-12 neurons per group). Nevertheless, none of the intrinsic properties differed significantly between TED mice and controls in either condition. Taken together with the morphological analyses, the data point to an increase in the extrinsic excitatory connections (via mature spines) on D1R SPNs, rather than a change in intrinsic excitability, as the mechanism that supports normal motor function in TED (and VGLUT3 KO) mice following dopamine depletion.

### Enhanced activation of dopamine 1 receptors is required for the preserved morphology of D1R SPNs and motor behavior after dopamine depletion

Our data show that a transient daily increase in midbrain dopamine neuron activity during dorsal striatal dopamine depletion preserves the density of mature spines as well as the complexity and length of dendritic arbors on D1R, but not D2R SPNs. The selectivity of this effect for D1R SPNs suggests a potential role for enhanced activation of D1 receptors on these neurons. To test this directly, we implanted cannulas bilaterally in the dorsal striatum to deliver the highly selective D1 receptor antagonist, SCH23390, to VGLUT3 KO mice and to deliver, as a control, saline to both VGLUT3 KO and WT cohorts (Figure 4A-C). To establish the dose of antagonist required to reduce striatal D1 receptor activity in VGLUT3 KO mice to WT levels, we measured locomotor activity (Figure 4C). The antagonist (30 ug) was delivered once per day prior to the onset of the active cycle for 14 days and the 6-OHDA (6 µg) was injected unilaterally into the dorsal striatum on day 7 (Figure 4A). Paw contacts in the cylinder were measured at the peak of the inactive cycle both before and one week after 6-OHDA injection. Prior to depletion, all animal groups showed normal bilateral paw contacts, as expected (N=4-7 mice per group) (Figure 4D). After depletion, control VGLUT3 KO mice infused with saline showed normal bilateral paw contacts (N=4 mice), while KO mice infused with the D1 receptor antagonist showed ipsilateral paw dominance (ipsilateral vs contralateral p<0.0001, N=7 mice), similar to depleted WT mice infused with saline (ipsilateral vs contralateral p<0.0001, N=5 mice) (Figure 4D). These data provide strong support for the idea that enhanced activation of D1 receptors during induction of the Parkinsonian state has a required role in the long-term preservation of motor function.

**Figure 4.**
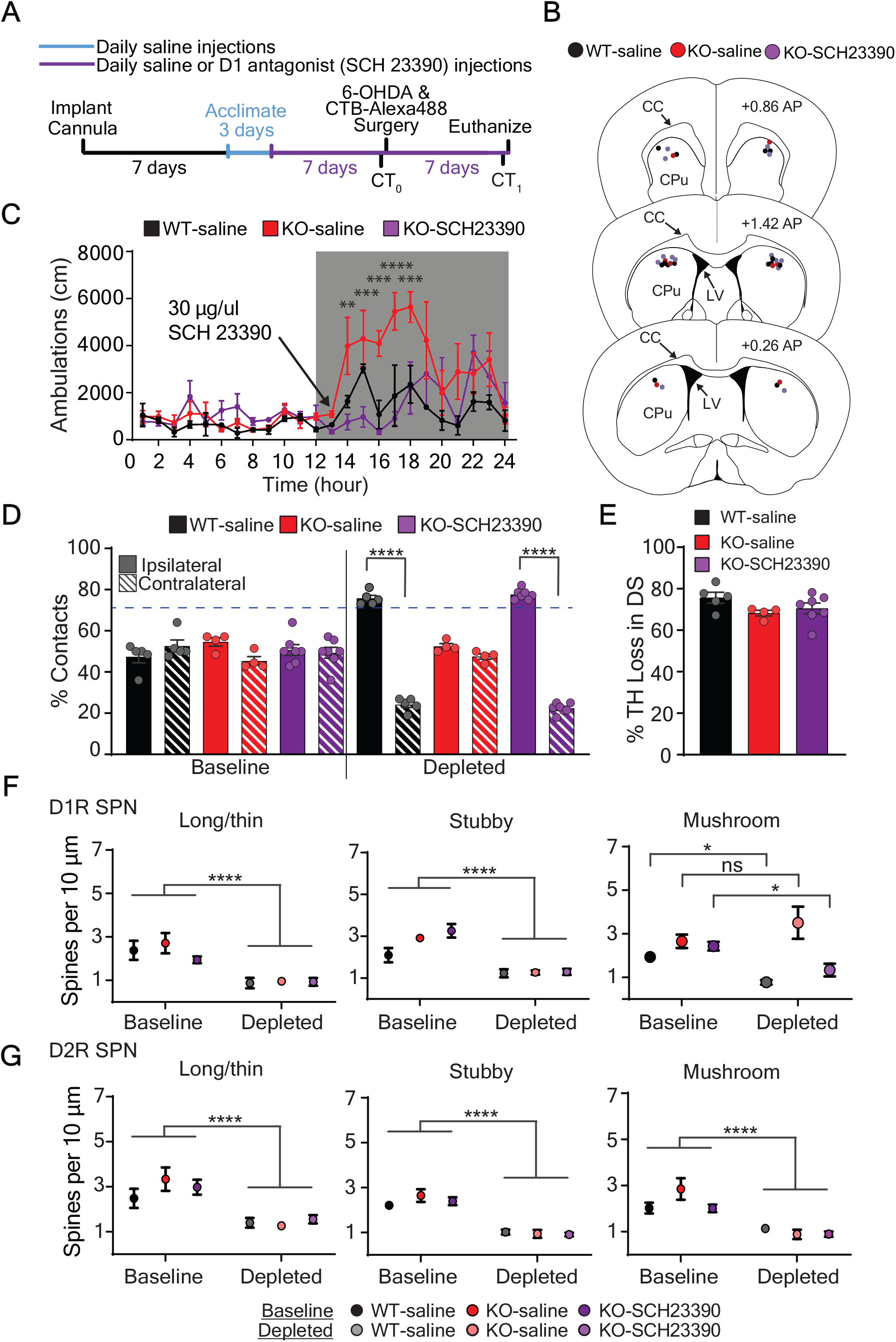
Elevated striatal dopamine signaling through D1 receptors during dorsal striatal dopamine depletion is required for preserved D1R SPN morphology and motor behavior. **(A)** Experimental paradigm for testing the role of enhanced striatal D1 receptor signaling on motor behavior and SPN morphology in a model of PD. CT=cylinder test. **(B)** Locations of bilateral cannula placement in dorsal striatum of the three animal groups. CC=corpus callosum CPu= caudate putamen, LV= lateral ventricle, AP= anterior posterior **(C)** The dose of D1 receptor antagonist (SCH23390, 30 µg in 1 ul) delivered directly into the dorsal striatum of VGLUT3 KO mice was optimized to suppress locomotor activity to WT levels. Significant interaction by 2-way ANOVA (*animal group* x *hour*) with Bonferroni post-hoc analysis. **(D)** One week after 6-OHDA injection, VGLUT3 KO mice infused with SCH23390 (purple striped bar) developed ipsilateral paw dominance similar to control WT mice infused with saline (black striped bar), while control VGLUT3 KO mice infused with saline (red striped bar) continued to show bilateral paw reaches as expected. Significant interaction by 2-way ANOVA (*animal group x condition*) with Bonferroni post-hoc analyses. **(E)** Percent decrease in dorsal striatal TH immunoreactivity one week after 6-OHDA injection is similar across all three animal groups, Kruskal-Wallis test. **(F)** At baseline, D1R SPN spine densities were similar between animal groups. After dopamine depletion, mature (mushroom) spine densities of VGLUT3 KO (D1 antagonist infused) were decreased similarly to the WT (saline infused), while control VGLUT3 KO (saline infused) showed the expected preserved mushroom spine densities. Significant interaction by 2-way ANOVA (*animal group x condition*) with Bonferroni post-hoc analyses. **(G)** Baseline D2R SPN spine densities were similar between animal groups. After depletion, spine densities were significantly decreased across animal groups. All measurements during inactive cycle. No interaction by 2-way ANOVA (*animal group x condition*). Significant main effects of condition. *p<0.05, **p<0.01, ***p<0.005, ****p<0.0001, ns = not significant.

Based on the work shown here, our model for the mechanism that allows normal motor function after dopamine depletion is the preservation of D1R SPN morphology. Therefore, we also measured mature spine density of D1R SPNs in the different treatment groups. After depletion as expected, the saline infused VGLUT3 KO mice showed preserved mature spine density on D1R SPNs (N=3-4 mice per group, n=6-9 dendrites per group) (Figure 4F). In contrast, antagonist infused VGLUT3 KO mice showed a decrease in mature spine density on D1R SPNs (baseline vs depleted p<0.05, N=4 mice per group, n=5-9 dendrites per group) that was similar to the decrease observed in saline infused WT mice (baseline vs depletion p<0.05; N=3 mice per group, n=6-9 dendrites per group) (Figure 4F). As expected, spine densities measured on both types of SPNs at baseline (during the inactive cycle) were the same for animal groups. Additionally, the spine densities measured on D2R SPNs following dopamine depletion were significantly decreased for animal groups (depleted vs non-depleted p<0.001, N=3-4 mice per group, n=6-9 dendrites per group) (Figure 4G). Importantly, the extent of dorsal striatal dopamine depletion was similar between all animal groups (N=3-7 mice per group) (Figure 4E).

## DISCUSSION

We report here the induction of a long-term form of striatal plasticity that supports normal paw reaching behavior following dopamine depletion. This form of plasticity is induced by a temporary, transient, daily increase in midbrain dopamine neuron activity and requires the enhanced activation of striatal D1 receptors. The plasticity manifests as a preservation of D1R, but not D2R SPN dendritic architecture. In contrast, the intrinsic excitability of the SPNs were similar across all mouse groups following depletion, and thus is not likely a contributing factor to the normal motor behavior. Taken together, the data suggest that the normal motor behavior of VGLUT3 KO and TED mice after dopamine depletion is due to a preservation of the excitatory synaptic connectivity onto D1R SPNs, which is consistent with the theory of a “restored” balance in the output pathways of the striatum (i.e. direct over indirect pathway output facilitates movement). The motor cortex is the likely source of excitatory inputs that maintain normal motor function (Ding et al., 2008), though plasticity of thalamic inputs has also been shown to affect motor function in PD (Parker et al., 2016). Future studies to functionally map synaptic level changes in cortical and thalamic input to the dorsal striatum in dopamine depleted VGLUT3 KO and TED mice, will provide further insight into the anatomical underpinnings of this remarkable and potentially therapeutically beneficial form of long-term striatal plasticity.

We used a model of PD that limits dopamine depletion to the nigrostriatal pathway, thus better mimicking the loss of dopamine that occurs primarily within the putamen in human PD, and particularly within the early stages (Boix et al., 2015). More commonly used 6-OHDA models target the median forebrain bundle, producing lesions of the midbrain dopamine system that are even more widespread than has been observed in late-stage human PD (Boix *et al*., 2015). Nevertheless, confirmation of these results using other models that similarly lead to a primarily nigrostriatal degeneration and that also show more of the pathologies (e.g. alpha-synuclein inclusions or genetic models) and/or mimic a slower degeneration as seen in subsets of PD patients, will provide important additional insight into the clinical utility of the plasticity mechanism identified here.

Next steps in the study of the plasticity will be to understand the transmitters and downstream intracellular signaling pathways that are involved in the preservation of the dendritic architecture beyond dopamine and D1 receptors. Dopamine neurons also release GABA, glutamate, and neuropeptides (Tritsch et al., 2012; Trudeau et al., 2014). Enhanced release of these neurotransmitters in VGLUT3 KOs and in the TED mice may also have a role. Because the plasticity was specific to D1R SPNs, we examined the requirement for enhanced D1 receptor activity. While there were no obvious changes in D2R SPNs in this model, D2 receptors located on dopamine or cortical terminals may nevertheless contribute as well.

Therapeutic strategies administered in the early stages of PD have the potential to delay the onset or slow the progression of the motor symptoms. However, preclinical studies of PD tend to focus on introducing interventions at time points after dopamine depletion is complete (late stage). Moreover, because long-term use of dopamine replacement therapy, the most common and effective treatment for PD symptoms, can lead to debilitating dyskinesias or cease working, the treatment is often not administered until the late stages of the disease. Data presented here provide a strong rationale to shift the therapeutic paradigm to an earlier stage and to be able to administer the memetics short-term.

Interestingly, exercise which is employed as an adjunct treatment for PD because of its neuroprotective and restorative effects (Palasz et al., 2019), elevates striatal dopamine and promotes spine enhancement within the mammalian brain (Hou et al., 2017), including on D1R and D2R SPNs in PD (Toy et al., 2014). The spine enhancement may be mediated though neurotrophins, which are secreted from both cortico- and nigrostriatal afferents (Baquet et al., 2004; Baydyuk and Xu, 2014; Conner et al., 1997). Neurotrophins play a critical role in neuronal growth, maintenance, and plasticity (Chao, 2003; Park and Poo, 2013), and have been shown to be neuroprotective in PD animals models (Nagahara and Tuszynski, 2011). Mechanisms similar the ones we report here thus may underlie the benefits of exercise in PD patients.

Additional potential strategies to induce this striatal mechanism at early stages of the disease for therapeutic purposes include identifying and manipulating the neurons that give rise to the hyperdopaminergic phenotype in the VGLUT3 KO mice. Striatal cholinergic interneurons are poised to modulate dopamine levels, but deletion of VGLUT3 from these interneurons did not produce hyperlocomoter activity during any period of the circadian cycle (Divito *et al*., 2015). Lastly, the dopamine-mediated plasticity described here may have implications for indications beyond motor function, for example, mood and cognitive symptoms that also result from midbrain dopamine deficiency and are commonly co-morbid in PD (Goldman et al., 2018; Marras and Chaudhuri, 2016).

## ACKNOWLEDGMENTS

We thank Dr. Myung-chul Noh for help with electrophysiological recordings and comments on the manuscript; Drs. David Ferreira and Cynthia Arokiaraj for experimental design input and comments on the manuscript; Dr. Aryn Gittis for technical advice; Drs. Bryan Hooks and Yan Dong for technical advice and comments on the manuscript; Cole Boillat, Daniel Butz, Jenna Nemschoff, Zane Hamden, Amy Liu, Maximillion Junge, Tulika Malik, Nikhill Khongbantabam, Thalia, Petroulakis, Randi Wilk, and Gopika Pillai for counting spines and scoring behavior; Jeremy Gedeon for technical assistance; Sean Paul Williams and Kelly Corrigan for mouse colony maintenance; Suh Jin Lee and Kelly Corrigan for comments on the manuscript. Center for Biological Imaging and Simon Watkins for confocal microscope support; Suh Jin Lee for illustration in Figure S1D. Funding was T32NS086749 (J.C.B.), R01NS082650 (R.P.S.), DoD-Army: W81XWH-17-1-0386 (R.P.S.), and a UPBI NeuroDiscovery Pilot Grant (R.P.S.).

## Author Contributions

R.P.S. designed the study; R.P.S. and J.B. designed experiments; J.B. performed the experiments, J.B and R.P.S analyzed and interpreted the data; J.B. and R.P.S. wrote the paper.

## STAR METHODS

## LEAD CONTACT AND RESOURCE AVAILABILITY

Further information and requests for reagents and resources should be directed and will be fulfilled by Rebecca Seal.

Any raw data obtained for this study are available from the lead contact upon request.

The study did not generate any code.

Any additional information required to reanalyze the data reported here is available from the lead contact upon request.

### Experimental Animal and subject details

Animals were housed in micro-isolator cages on a reverse 12h active/inactive cycle (10 A.M. lights off, 10 P.M. lights on). All animals had *ad libitum* access to food and water and were treated in compliance with Institutional Animal Care and Use Committee for the University of Pittsburgh. All efforts were made to minimize the number of animals used and to avoid pain and discomfort. In each experiment, approximately equal numbers of males and females were used. *Slc17a8*^-/-^mice (also referred to as VGLUT3^-/-^or VGLUT3 KO mice) were backcrossed at least 10 generations to C57BL/6(5). DAT^cre^ (RRID: IMSR_JAX:020080) and D1^dt^ (RRID: IMSR_JAX:016204) were obtained from Jackson laboratories. For eDREADD experiments DAT^cre^ mice were utilized. For electrophysiological eDREADD experiments, DAT^cre^ and D1^dt^ mice were crossed to obtain DAT^cre^; D1^dt^ mice. When testing the duration of normal motor behavior in VGLUT3 KO mice, following DA depletion, mice began the experiment between the age of 3-5 months and ended the experiment between 8-10 months. For all remaining experiments, mice were 3-6 months in age.

### Animal Groups and Experimental Approach

Experiment 1: SPN spine morphology in a dopamine depleted state. VGLUT3 KO and WT mice were used. Experiment 2: Effects of transient elevated dopamine on motor behavior, spine dynamics, and electrophysiology of SPNs in normal and dopamine depleted states. DAT^cre^; D1^dt^ mice were used for electrophysiology experiments and DAT^cre^ mice were used for the remaining experiments. Mice in these experiments were randomly divided into three groups: 1) mCherry+CNO, 2) eDREADD+Saline, or 3) eDREADD+CNO. At 3 months of age mice were injected with the eDREADD or mCherry virus bilaterally into the SNpc. At least three weeks following viral injection eDREADD and mCherry animals were either given saline or CNO one hour before the onset of the active cycle for 5 days. These animals then underwent baseline behavioral testing. The injections continued daily for 9 days following dopamine depletion via 6-OHDA. Animals were assessed behaviorally, one-week following depletion. Animals were euthanized during the inactive and active cycle and assessed for spine and dendrite morphology and electrophysiology.

### Behavior

Light conditions and temperature were kept at ambient levels (∼32 lux, 75°C). For home-cage locomotor activity, mice were placed in rat micro-isolator cages inside a photobeam monitoring system (Kinder Scientific) for 72 hours with food and water *ad libitum*. For the *cylinder test*, mice were placed in a clear glass cylinder ∼8 cm in diameter with a clear acrylic sheet on top. The video camera was placed overhead and recorded one 5-minute video per animal per condition. The number of full weight-bearing forepaw contacts (identified as the majority of the forepaw coming in contact with the cylinder) against the cylinder wall was recorded and expressed as total percentage of contacts. Paw dominance is equal to or greater than 70% ipsilateral paw contacts in the cylinder test. Animals were tested 3-4 hours before and after the start of the active cycle on separate days. Videos with less than 7 total rears were excluded from analysis. All videos were scored by two trained observes unaware of the treatment groups. Their scores were compiled and averaged together.

### Stereotaxic Surgery

All surgical procedures were performed using aseptic technique. Mice were anesthetized with a 3% induction/1-3% maintenance of isoflurane/100% oxygen mixture. Specific details of surgery can be found in Divito et al 2015. Briefly, for eDREADD injections, mice were either given 1 µL of AAV8-hSyn-DIO-hM4d(Gi)-mCherry (**RRID**: Addgene_44362) or AAV8-hSyn-DIO-mCherry (**RRID**: Addgene_50459) bilaterally into the SNpC (in mm) - 3.29 AP, ±1.30 ML, and -4.40 DV. For 6-hydroxydopamine hydrochloride (6-OHDA) injections, mice were given unilateral partial dorsal striatal lesions to assess Parkinsonian motor behavior. Stereotaxic coordinates used for the unilateral injection of 6-OHDA (6 µg, Sigma) or saline were (in mm) +0.85 AP, -1.75 ML, and -2.75 DV. Injections were administered at 0.25µL/min with three-minute incubation periods following needle placement and solution injection. Immediately following surgery, mice were sutured and administered ketoprofen (Ketofen 5 mg/kg, Zoetis) and allowed to recover for one week.

Mice also received IP injections of 0.9% saline (0.5mL) 1-2x daily for 6 days after surgery. Mouse weight was monitored and any that lost >20% of their presurgical weight were excluded. No animals were lost or euthanized after surgery due to surgical complications. 6-OHDA injected animals with less than 70% decrease in tyrosine hydroxylase (TH) staining across the rostral-caudal extent of the dorsal striatum were excluded from analysis. Baseline behavior tests were performed at least three-weeks following eDREADD injections and post-depleted behaviors were assessed one-week following 6-OHDA depletion. Brains were either harvested for spine analysis or electrophysiology (see below).

### Cannulation Surgery and Infusion Procedures

Mice were anesthetized with a 3% induction/1-3% maintenance of isoflurane/100% oxygen mixture. An incision was made in the midline of the skull and 2 small burr holes were drilled through the skull at stereotaxic coordinates +0.85 AP and -/+1.75 ML. Low profile 26 Ga, 1.75 mm guide cannulas (Plastics One 42593), containing a 33 Ga, 2.75mm internal dummy cannula (Plastics One 42597) were implanted into these coordinates at a depth of -2.75 DV and secured with GC FujiCem 2 glass ionomer cement (Fisher, 5081153). The wound was sutured closed if any remaining bone or musculature were exposed. Ketoprofen (Ketofen 5 mg/kg, Zoetis) was administered immediately following surgery and the mice allowed to recover for one week. Mice also received IP injections of 0.9% saline (0.5mL) 1-2x daily for 6 days after surgery. Mice were singly housed following surgical procedures to avoid accidental loss of cannulas. Mouse weight was monitored and any that lost >20% of their presurgical weight were excluded. To infuse saline or D1R antagonist (Tocris, SCH-23390) bilaterally into the striatum, mice were first pinned down with a cloth, had their internal dummy cannulas removed, and placed the internal infusion cannula (Plastics One, 42595) into the right and left guide cannula. 1µL of saline or D1R antagonist (SCH-23390, TOCRIS) were infused at a rate of 0.25µl/min into each hemisphere while the mice moved freely. Three minutes following infusion, the mouse was again pinned down, the infusion cannulas removed, the internal dummy cannula inserted, and the mouse was placed back into the home cage. Prior to behavioral testing, all mice were acclimated to infusion procedures with 3 days of saline injections.

### DiI labeling and Immunohistochemistry

To analyze spine morphology, mice were euthanized with an overdose of ketamine/xylazine and intracardially perfused with 25 mL PBS (in mm): 7.95 NaCl, 0.20 KCl, 1.425 Na_2_HPO_4_, 0.27 KH_2_PO_4_, followed by 50mL of cold PBS with 2.5% paraformaldehyde in PBS (para). Brains were dissected and postfixed for one hour in 2.5% para. Brains were then sectioned at 100µm on a vibratome (LEICA VT1000S) and every other section was wet mounted onto a slide. Small DiI (Thermofisher, D3911) crystals were then placed on the dorsal striatum via glass micropipette. Sections were incubated in PBS at 4°C for two days and subsequently fixed with 4% para for one hour at room temperature. The slides were then coverslipped and stored at 4°C until imaged on a confocal. The remaining slices were fixed in 4% para before being stained for TH immunoreactivity. Slices were washed (3×5 minutes) in PBS and incubated in 1:1000 Rabbit α-tyrosine hydroxylase (RRID: AB_390204) containing 0.4%Triton X-100 and 5% normal donkey serum in PBS for 48 hours at room temperature. After washing, these slices were then incubated in 1:500 Alexa Fluor 488 conjugated donkey α-rabbit (RRID: AB_2313584) containing 0.4%Triton X-100 for two hours at room temperature and then mounted with DAPI Fluromount-G (southern Biotech, 0100-20). For electrophysiology experiments, a 1:500 Alexa Fluor 647 streptavidin (RRID: AB_2341101) was added to the secondary solution for neurobiotin labeling. Labeled neurons were imaged on a confocal at 40x with NIS-elements (RRID: SCR_014329) software, then traced and analyzed for Sholl intersections using Simple Neurite Tracer in Fiji-ImageJ (RRID: SCR_002285) software.

### Electrophysiology

Coronal sections (300 µm thickness) were prepared from brains of mice 3-6 months in age of both sexes and euthanized before the onset of the active cycle (10 am). Slices were cut in ice-cold, carbogenated, N-methyl-D-glucamine-HEPES solution containing the following (in mM): 93 N-methyl-D-glucamine, 2.5 KCl, 1.2 NaH_2_PO_4_, 30 NaHCO_3_, 20 HEPES, 25 glucose,10 MgSO_4_, 0.5 CaCl_2_, 5 sodium ascorbate, 2 thiourea, and 3 sodium pyruvate, pH 7.3. Slices were allowed to recover for 15 minutes at 33°C in this solution before being held at room temperature for 1 hour in carbogenated artificial cerebral spinal fluid (ACSF) as follows (in mM): 125 NaCl, 26 NaHCO_3_, 1.25 NaH_2_PO_4_, 2.5 KCl, 12.5 glucose, 1 MgCl_2_, and 2 CaCl_2_. Recordings were made at 33°C using carbogenated ASCF. Whole-cell patch-clamp recordings were performed using borosilicate pipettes (4-6 MΩ resistance) filled with an internal solution containing the following (in mM): 130 KMeSO_3_, 10 NaCl, 2 MgCl_2_, 0.16 CaCl_2_, 0.5 EGTA, 10 HEPES, 2 Mg-ATP, and 0.3 Na-GTP with 0.2% Neurobiotin (RRID: AB_2336606), pH 7.3. Cells in which series resistance changed >20% over the course of the recording were excluded from analysis. Data were collected and analyzed using pClamp-10 (RRID: SCR_011323) software.

### Statistics

The Kruskal-Wallis Test was used to analyze the percent decrease in TH immunoreactivity in the dorsal striatum. A mixed effect model with repeated measures was used to analyze depleted animals over the course of 20 weeks. For the remaining analyses, we used a two-way ANOVA, repeated measures analysis (*condition vs genotype or condition vs mouse type*). If no significant interaction was found, we reported the main effect. If a significant interaction was found, we used *post-hoc* comparisons with a Bonferroni correction. All data were measured for normality against a Gaussian distribution, using both Shapirp-Wilk and Kolmogorov-Smirnov tests (alpha=0.05). Significance was determined at p<0.05. All data are represented as mean ± SEM. All analyses were performed in GraphPad Prism 8 (RRID: SCR_002798) software. ^*^p<0.05, ^**^p<0.01, ^***^p<0.001, ^****^p<0.0001

## SUPPLEMENTAL INFORMATION

Document S1. Figures S1-S3.

## Supplemental Material

### Supplemental Figures and Legends

**Figure S1.**
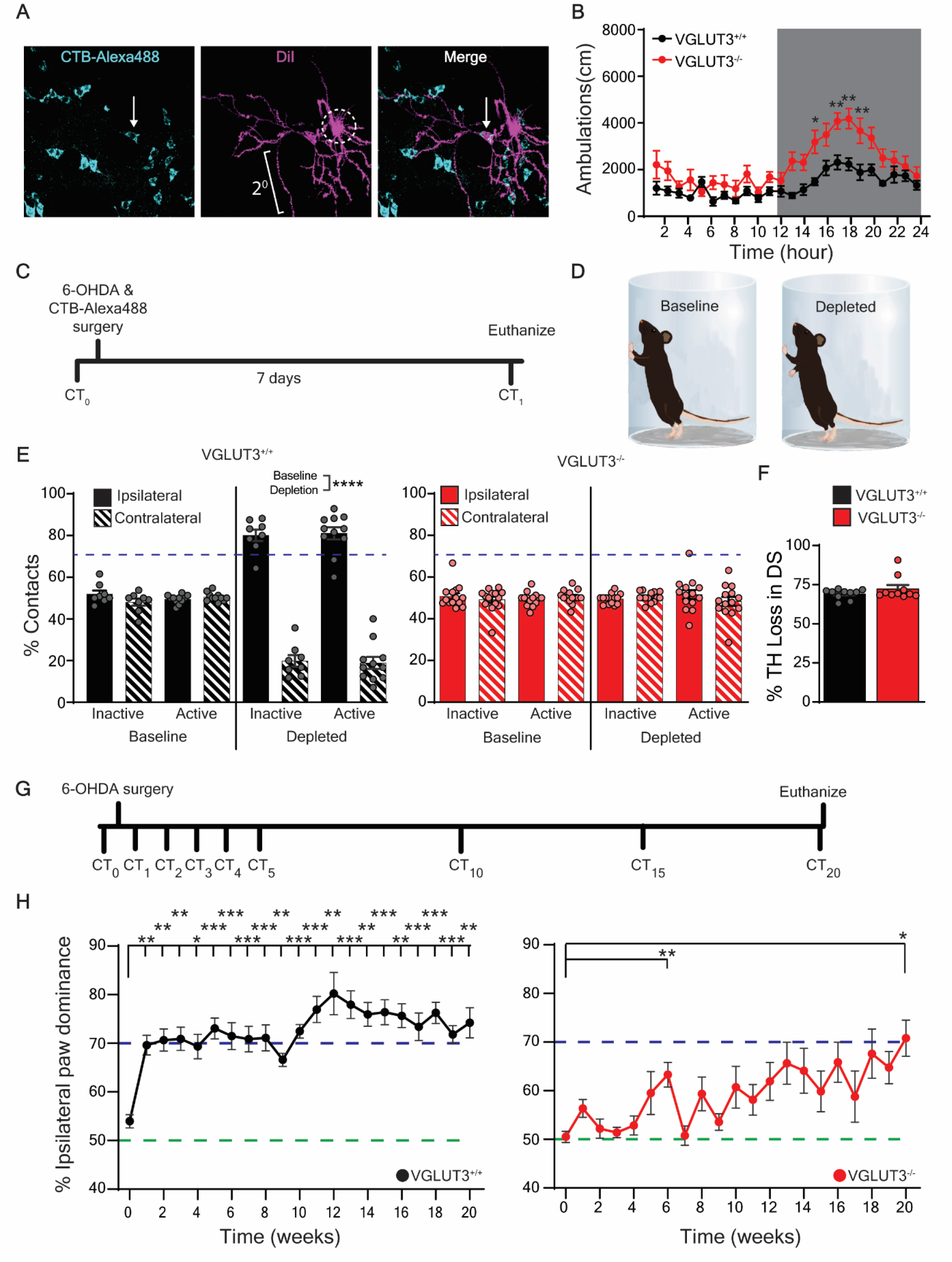
Preserved motor function of VGLUT3 KO in a model of PD. **(A)** D1R SPNs (white arrow, cyan) are back-labeled from SNpR by CTB-Alexa488 injection. DiI (magenta) crystal (white dotted outline) applied to acute striatal slices labels the membranes of SPNs for spine analysis of secondary dendrites (white 2°). **(B)** VGLUT3 KO mice are hyperlocomotive compared to WT mice during the waking cycle. Significant interaction by 2-way ANOVA *(genotype* x *time)* followed by Bonferroni *post-hoc* correction. **(C)** Experimental paradigm for testing changes in motor behavior and SPN morphology due to unilateral dorsal striatal 6-OHDA mediated dopamine depletion. CT= cylinder test. **(D)** Illustration of the cylinder test for motor behavior. Mice normally show bilateral paw reaches at baseline (left) and develop ipsilateral paw dominance after unilateral dopamine depletion (right). **(E)** Both VGLUT3 KO and WT mice show bilateral paw reaches at baseline. By one week after depletion, WT (black) mice have developed ipsilateral paw dominance, while VGLUT3 KO (red) continue to show bilateral paw reaches. Significant interaction by 2-way ANOVA (*genotype x condition*) followed by Bonferroni *post-hoc* correction. **(F)** Percent decrease in dorsal striatal tyrosine hydroxylase (TH) immunoreactivity after 6-OHDA injection is similar for KO and WT mice. N=10, Kruskal-Wallis. **(G)** Experimental timeline for a long-term depletion. CT= cylinder test, CT_0_ = CT at baseline, CT_x_ = x weeks after depletion. **(H)** Percent ipsilateral paw reaches become significant by one week after dopamine depletion in WT control mice and consistently significant by 20 weeks in VGLUT3 KO mice. Purple and green dotted lines represent ipsilateral paw dominance and no paw dominance, respectively. One-way ANOVA with Bonferroni *post-hoc* comparisons to baseline. N=11 WT, N=11 KO. *p<0.05, **p<0.01, ***p<0.005, ****<0.001. Related to Figure 1.

**Figure S2.**
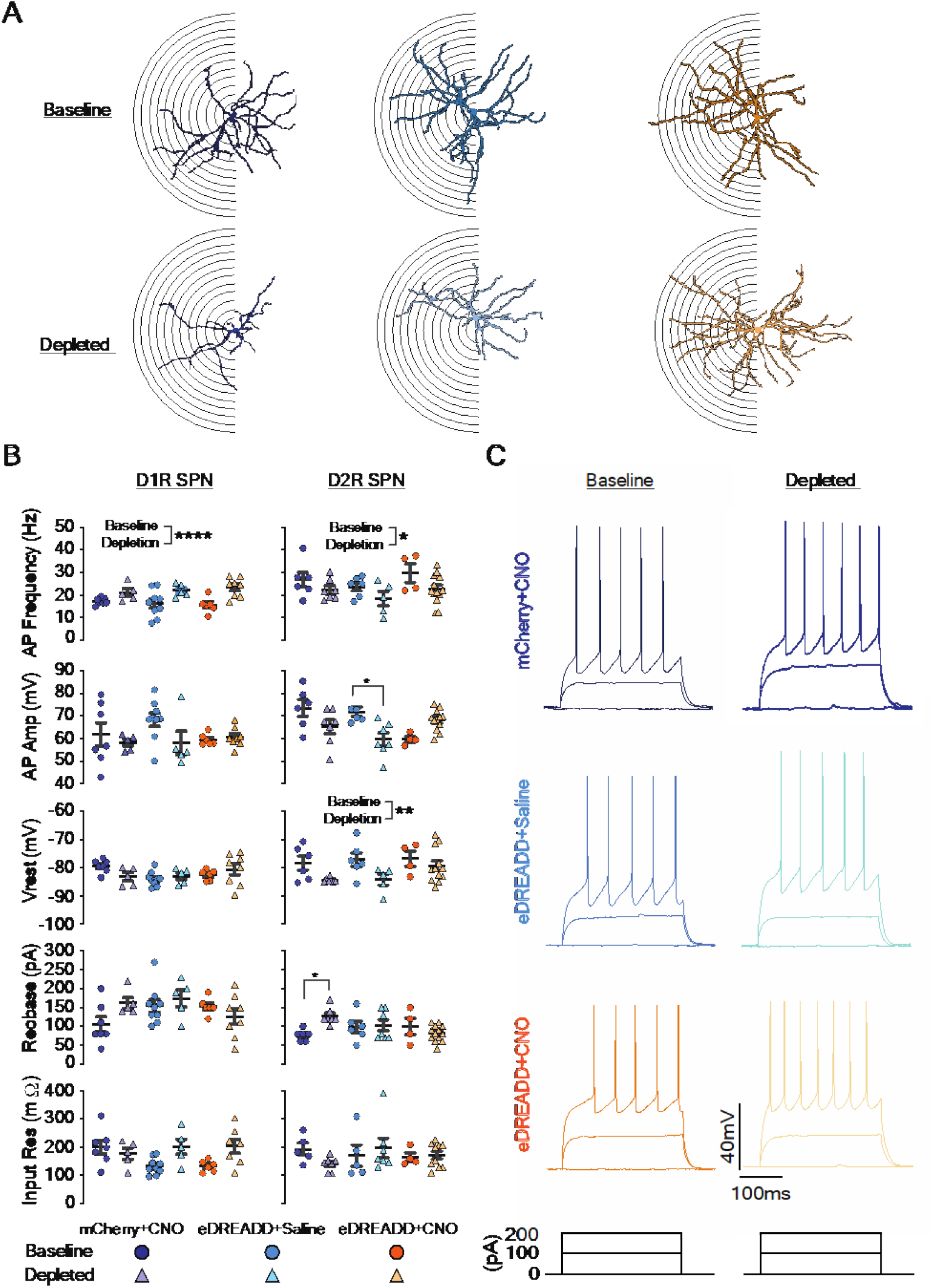
Sholl analysis depiction and intrinsic electrophysiological properties of SPNs in the eDREADD paradigm. **(A)** Representative image Sholl analysis for D1R SPNs at baseline (top) and after dopamine depletion (bottom). mChery+CNO (darkblue), eDREADD+Saline (light blue), and eDREADD+CNO (orange). **(B)** D1R SPNs (left) show an increase in action potential (AP) frequency in all three animal groups: mCherry+CNO (dark blue), eDREADD+Saline (light blue), and eDREADD+CNO (orange) one week after dopamine depletion. No changes were observed in AP amplitude, Vrest, rheobase, or input resistance. No significant interaction by 2-way ANOVA (*animal group* x *condition*). Significant main effect of condition. D2R SPNs (right) from all animal groups show a decrease in action potential frequency and Vrest following dopamine depletion. No significant interaction by 2-way ANOVA (*animal group* x *condition*). Significant main effect of condition. **C**, Example traces of D1R SPNs at baseline (right, top) and after depletion (left, top). Current injection paradigm (bottom). *p<0.05, **p<0.01, ****p<0.0001. Related to Figure 3.

**Figure S3.**
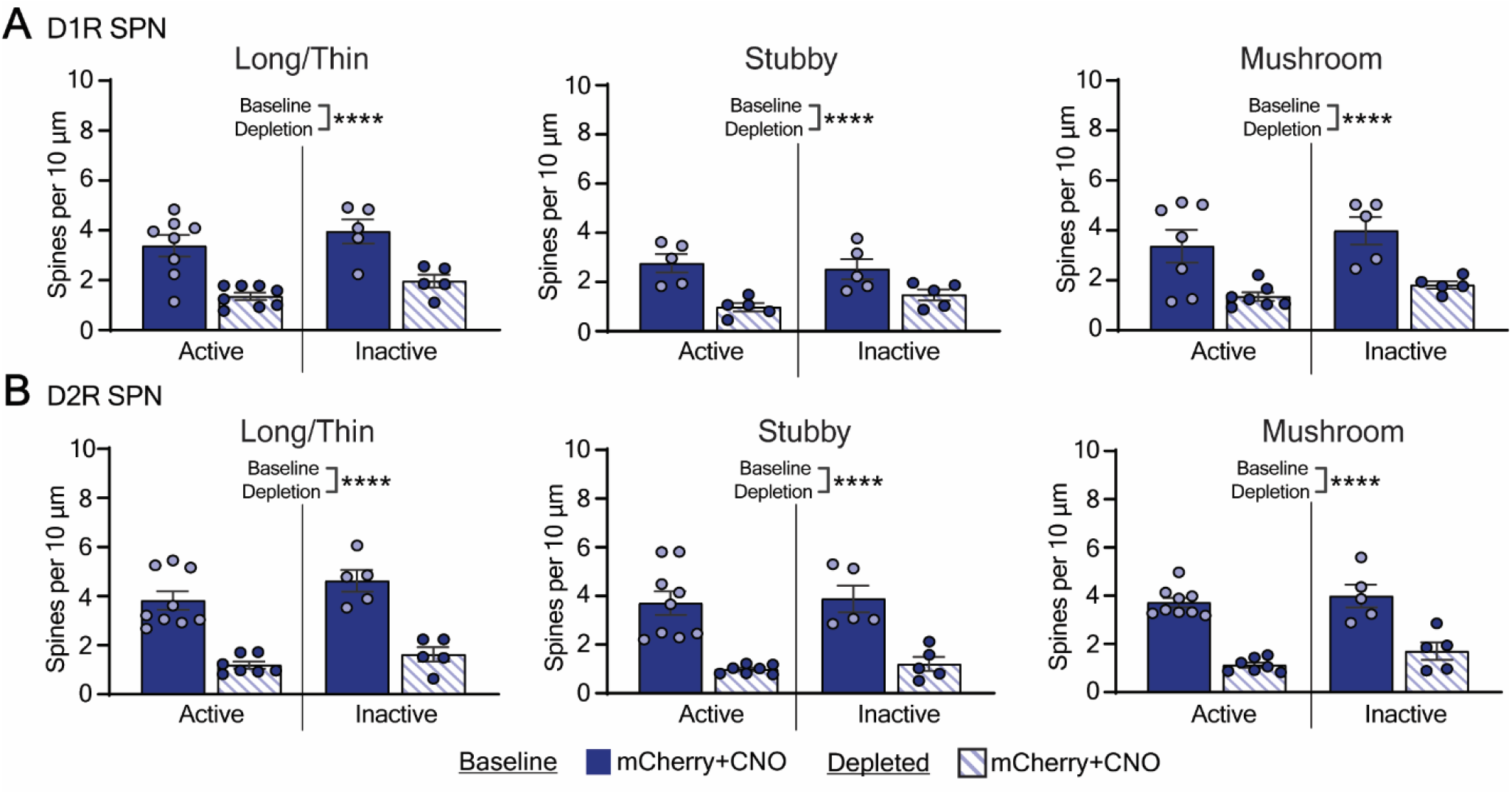
Changes in SPN spine densities in mCherry control mice observed after dorsal striatal dopamine depletion. **(A)** At baseline, long/thin, stubby, and mushroom D1R SPN spine density in mCherry+CNO mice (dark blue solid bars) is enhanced compared to depleted SPNs (striped blue bars) across the circadian cycle. No interaction by 2-way ANOVA (*animal group x condition*). Significant main effect for condition. **(B)** Depletion (striped blue bars) decreased D2R SPN spine densities (long/thin, stubby, mushroom) when compared with baseline (dark blue). No interaction by 2-way ANOVA (*animal group x condition*). Significant main effect for condition. ****p<0.001. Related to Figure 3.

## KEY RESOURCES TABLE

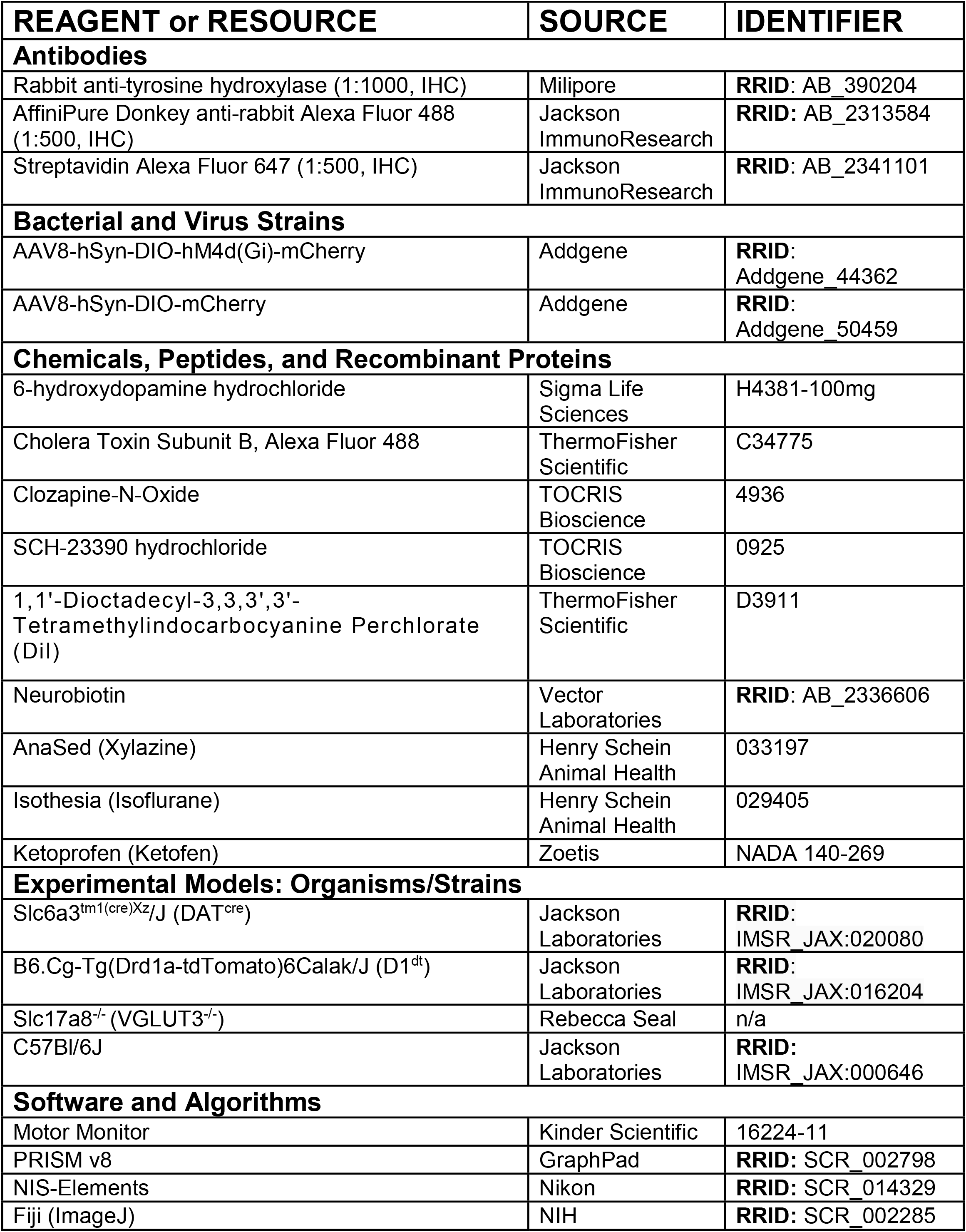
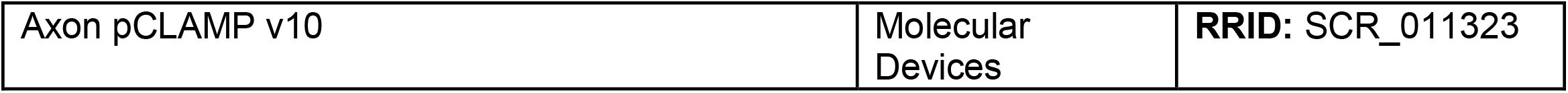

| REAGENT or RESOURCE | SOURCE | IDENTIFIER |
| --- | --- | --- |
| **Antibodies** |  |  |
| Rabbit anti-tyrosine hydroxylase (1:1000, IHC) | Milipore | RRID: AB_390204 |
| AffiniPure Donkey anti-rabbit Alexa Fluor 488 (1:500, IHC) | Jackson ImmunoResearch | RRID: AB_2313584 |
| Streptavidin Alexa Fluor 647 (1:500, IHC) | Jackson ImmunoResearch | RRID: AB_2341101 |
| **Bacterial and Virus Strains** |  |  |
| AAV8-hSyn-DIO-hM4d(Gi)-mCherry | Addgene | RRID: Addgene_44362 |
| AAV8-hSyn-DIO-mCherry | Addgene | RRID: Addgene_50459 |
| **Chemicals, Peptides, and Recombinant Proteins** |  |  |
| 6-hydroxydopamine hydrochloride | Sigma Life Sciences | H4381-100mg |
| Cholera Toxin Subunit B, Alexa Fluor 488 | ThermoFisher Scientific | C34775 |
| Clozapine-N-Oxide | TOCRIS Bioscience | 4936 |
| SCH-23390 hydrochloride | TOCRIS Bioscience | 0925 |
| 1,1'-Diocadecyl-3,3,3',3'-Tetramethylindocarbocyanine Perchlorate (DiI) | ThermoFisher Scientific | D3911 |
| Neurobiotin | Vector Laboratories | RRID: AB_2336606 |
| AnaSed (Xylazine) | Henry Schein Animal Health | 033197 |
| Isothesia (Isoflurane) | Henry Schein Animal Health | 029405 |
| Ketoprofen (Ketofen) | Zoetis | NADA 140-269 |
| **Experimental Models: Organisms/Strains** |  |  |
| Slc6a3<sup>tm1(cre)Xz/J</sup> (DAT<sup>cre</sup>) | Jackson Laboratories | RRID: IMSR_JAX:020080 |
| B6.Cg-Tg(Drd1a-tdTomato)6Calak/J (D1<sup>dt</sup>) | Jackson Laboratories | RRID: IMSR_JAX:016204 |
| Slc17a8<sup>-/-</sup> (VGLUT3<sup>-/-</sup>) | Rebecca Seal | n/a |
| C57Bl/6J | Jackson Laboratories | RRID: IMSR_JAX:000646 |
| **Software and Algorithms** |  |  |
| Motor Monitor | Kinder Scientific | 16224-11 |
| PRISM v8 | GraphPad | RRID: SCR_002798 |
| NIS-Elements | Nikon | RRID: SCR_014329 |
| Fiji (ImageJ) | NIH | RRID: SCR_002285 |
| Axon pCLAMP v10 | Molecular<br>Devices | **RRID:** SCR_011323 |

## REFERENCES

Albin, R.L., Young, A.B., and Penney, J.B. (1989). The functional anatomy of basal ganglia disorders. 12, 336–375.

Armbruster, B.N., Li, X., Pausch, M.H., Herlitze, S., and Roth, B.L. (2007). Evolving the lock to fit the key to create a family of G protein -coupled receptors potently activated by an inert ligand. PNAS 104, 5163–5168.

Baquet, Z.C., Gorski, J.A., and Jones, K.R. (2004). Early striatal dendrite deficits followed by neuron loss with advanced age in the absence of anterograde cortical brain-derived neurotrophic factor. J Neurosci 24, 4250–4258. 10.1523/JNEUROSCI.3920-03.2004.

Baydyuk, M., and Xu, B. (2014). BDNF signaling and survival of striatal neurons. Front Cell Neurosci 8, 254. 10.3389/fncel.2014.00254.

Boix, J., Padel, T., and Paul, G. (2015). A partial lesion model of Parkinson’s disease in mice--characterization of a 6-OHDA-induced medial forebrain bundle lesion. Behav Brain Res 284, 196–206. 10.1016/j.bbr.2015.01.053.

Chao, M.V. (2003). Neurotrophins and their receptors: a convergence point for many signalling pathways. Nat Rev Neurosci 4, 299–309. 10.1038/nrn1078.

Conner, J.M., Lauterborn, J.C., Yan, Q., Gall, C.M., and Varon, S. (1997). Distribution of Brain-Derived Neurotrophic Factor (BDNF) Protein and mRNA in the Normal Adult Rat CNS: Evidence for Anterograde Axonal Transport. J Neurosci 17, 2295–2313.

Day, M., Wang, Z., Ding, J., An, X., Ingham, C.A., Shering, A.F., Wokosin, D., Ilijic, E., Sun, Z., Sampson, A.R., et al. (2006). Selective elimination of glutamatergic synapses on striatopallidal neurons in Parkinson disease models. Nat Neurosci 9, 251–259. 10.1038/nn1632.

Ding, J., Peterson, J.D., and Surmeier, D.J. (2008). Corticostriatal and thalamostriatal synapses have distinctive properties. J Neurosci 28, 6483–6492. 10.1523/JNEUROSCI.0435-08.2008.

Divito, C.B., Steece-Collier, K., Case, D.T., Williams, S.P., Stancati, J.A., Zhi, L., Rubio, M.E., Sortwell, C.E., Collier, T.J., Sulzer, D., et al. (2015). Loss of VGLUT3 Produces Circadian-Dependent Hyperdopaminergia and Ameliorates Motor Dysfunction and l-Dopa-Mediated Dyskinesias in a Model of Parkinson’s Disease. J Neurosci 35, 14983–14999. 10.1523/JNEUROSCI.2124-15.2015.

Fearnley, J.M.L. A.J. (1991). Ageing and Parkinson’s disease: substantia nigra regional selectivity. Brain : a journal of neurology 114, 2283–2301.

Fieblinger, T., Graves, S.M., Sebel, L.E., Alcacer, C., Plotkin, J.L., Gertler, T.S., Chan, C.S., Heiman, M., Greengard, P., Cenci, M.A., and Surmeier, D.J. (2014). Cell type-specific plasticity of striatal projection neurons in parkinsonism and L-DOPA-induced dyskinesia. Nat Commun 5, 5316. 10.1038/ncomms6316.

Gagnon, D., Petryszyn, S., Sanchez, M.G., Bories, C., Beaulieu, J.M., De Koninck, Y., Parent, A., and Parent, M. (2017). Striatal Neurons Expressing D1 and D2 Receptors are Morphologically Distinct and Differently Affected by Dopamine Denervation in Mice. Sci Rep 7, 41432. 10.1038/srep41432.

Goldman, J.G., Vernaleo, B.A., Camicioli, R., Dahodwala, N., Dobkin, R.D., Ellis, T., Galvin, J.E., Marras, C., Edwards, J., Fields, J., et al. (2018). Cognitive impairment in Parkinson’s disease: a report from a multidisciplinary symposium on unmet needs and future directions to maintain cognitive health. NPJ Parkinsons Dis 4, 19. 10.1038/s41531-018-0055-3.

Hou, L., Chen, W., Liu, X., Qiao, D., and Zhou, F.M. (2017). Exercise-Induced Neuroprotection of the Nigrostriatal Dopamine System in Parkinson’s Disease. Front Aging Neurosci 9, 358. 10.3389/fnagi.2017.00358.

Huang, Y.H., Lin, Y., Mu, P., Lee, B.R., Brown, T.E., Wayman, G., Marie, H., Liu, W., Yan, Z., Sorg, B.A., et al. (2009). In vivo cocaine experience generates silent synapses. Neuron 63, 40–47. 10.1016/j.neuron.2009.06.007.

Lanciego, J.L., Luquin, N., and Obeso, J.A. (2012). Functional neuroanatomy of the basal ganglia. Cold Spring Harb Perspect Med 2, a009621. 10.1101/cshperspect.a009621.

Lee, B.R., and Dong, Y. (2011). Cocaine-induced metaplasticity in the nucleus accumbens: silent synapse and beyond. Neuropharmacology 61, 1060–1069. 10.1016/j.neuropharm.2010.12.033.

Lee, B.R., Ma, Y.Y., Huang, Y.H., Wang, X., Otaka, M., Ishikawa, M., Neumann, P.A., Graziane, N.M., Brown, T.E., Suska, A., et al. (2013). Maturation of silent synapses in amygdala-accumbens projection contributes to incubation of cocaine craving. Nat Neurosci 16, 1644–1651. 10.1038/nn.3533.

Marras, C., and Chaudhuri, K.R. (2016). Nonmotor features of Parkinson’s disease subtypes. Mov Disord 31, 1095–1102. 10.1002/mds.26510.

McGregor, M.M., and Nelson, A.B. (2019). Circuit Mechanisms of Parkinson’s Disease. Neuron 101, 1042–1056. 10.1016/j.neuron.2019.03.004.

Nagahara, A.H., and Tuszynski, M.H. (2011). Potential therapeutic uses of BDNF in neurological and psychiatric disorders. Nat Rev Drug Discov 10, 209–219. 10.1038/nrd3366.

Palasz, E., Niewiadomski, W., Gasiorowska, A., Wysocka, A., Stepniewska, A., and Niewiadomska, G. (2019). Exercise-Induced Neuroprotection and Recovery of Motor Function in Animal Models of Parkinson’s Disease. Front Neurol 10, 1143. 10.3389/fneur.2019.01143.

Park, H., and Poo, M.M. (2013). Neurotrophin regulation of neural circuit development and function. Nat Rev Neurosci 14, 7–23. 10.1038/nrn3379.

Parker, P.R., Lalive, A.L., and Kreitzer, A.C. (2016). Pathway-Specific Remodeling of Thalamostriatal Synapses in Parkinsonian Mice. Neuron 89, 734–740. 10.1016/j.neuron.2015.12.038.

Smith, Y., Raju, D., Nanda, B., Pare, J.F., Galvan, A., and Wichmann, T. (2009). The thalamostriatal systems: anatomical and functional organization in normal and parkinsonian states. Brain Res Bull 78, 60–68. 10.1016/j.brainresbull.2008.08.015.

Smith, Y., and Villalba, R. (2008). Striatal and extrastriatal dopamine in the basal ganglia: an overview of its anatomical organization in normal and Parkinsonian brains. Mov Disord 23 Suppl 3, S534–547. 10.1002/mds.22027.

Smith, Y., Villalba, R.M., and Raju, D.V. (2009). triatal spine plasticity in Parkinson’s disease: pathological or not? Parkinsonism and Related Disorders 15S3, S156–S161.

Stephens, B., Mueller, A.J., Shering, A.F., Hood, S.H., Taggart, P., Arbuthnott, G.W., Bell, J.E., Kilford, L., Kingsbury, A.E., Daniel, S.E., and Ingham, C.A. (2005). Evidence of a breakdown of corticostriatal connections in Parkinson’s disease. Neuroscience 132, 741–754. 10.1016/j.neuroscience.2005.01.007.

Surmeier, D.J. (2007). Calcium, ageing, and neuronal vulnerability in Parkinson’s disease. The Lancet Neurology 6, 933–938. 10.1016/s1474-4422(07)70246-6.

Toy, W.A., Petzinger, G.M., Leyshon, B.J., Akopian, G.K., Walsh, J.P., Hoffman, M.V., Vuckovic, M.G., and Jakowec, M.W. (2014). Treadmill exercise reverses dendritic spine loss in direct and indirect striatal medium spiny neurons in the 1-methyl-4-phenyl-1,2,3,6-tetrahydropyridine (MPTP) mouse model of Parkinson’s disease. Neurobiol Dis 63, 201–209. 10.1016/j.nbd.2013.11.017.

Tritsch, N.X., Ding, J.B., and Sabatini, B.L. (2012). Dopaminergic neurons inhibit striatal output through non-canonical release of GABA. Nature 490, 262–266. 10.1038/nature11466.

Trudeau, L.E., Hnasko, T.S., Wallen-Mackenzie, A., Morales, M., Rayport, S., and Sulzer, D. (2014). The multilingual nature of dopamine neurons. Prog Brain Res 211, 141–164. 10.1016/B978-0-444-63425-2.00006-4.

Villalba, R.M., and Smith, Y. (2010). Striatal spine plasticity in Parkinson’s disease. Front Neuroanat 4, 133. 10.3389/fnana.2010.00133.

Villalba, R.M., and Smith, Y. (2018). Loss and remodeling of striatal dendritic spines in Parkinson’s disease: from homeostasis to maladaptive plasticity? J Neural Transm (Vienna) 125, 431–447. 10.1007/s00702-017-1735-6.

Zaja-Milatovic, S., Milatovic, D. Schantz, A.M., Zhang, J., Montine, K.S., Samii, A., Deutch, A. Y., and Montine, T. J. (2005). Dendritic Degeneration in Neostriatal Medium Spiny Neurons in Parkinson Disease. Neurology 64, 545–547.

